# MAP1A cleavage regulates TRIM46 transport to the axon in mature neurons

**DOI:** 10.64898/2026.06.27.735013

**Authors:** Macarena Pavez, Emma K. Gowing, Laura F. Gumy

**Affiliations:** Department of Anatomy, Faculty of Biomedical Sciences, University of Otago, Dunedin 9054, New Zealand

## Abstract

Polarised cargo transport is fundamental to neuronal function, ensuring the proper distribution of proteins and organelles between the cell body and axon. This process is tightly regulated at the proximal axon, where the enrichment of specific proteins organises the local cytoskeleton and directs intracellular trafficking. Although several microtubule-associated proteins (MAPs) have been implicated in axonal cargo transport during neuronal development, how protein localisation is maintained in mature neurons remains poorly understood. Here, we identify a previously unrecognised role for MAP1A in transporting TRIM46, a key organiser of the axonal microtubule cytoskeleton, to the axon of mature neurons, where it supports axon morphology. We show that MAP1A is enriched in the proximal axon and that its cleavage by Calpain-10 into heavy and light chains is required for TRIM46 localisation to this compartment. Mechanistically, the MAP1A light chain interacts with both TRIM46 and the tail domains of KIF3 motors, defining a transport complex that facilitates TRIM46 delivery to the proximal axon. Together, these findings establish proteolytic processing of a microtubule-associated protein as a mechanism regulating axonal protein distribution and neuronal polarity in mature neurons.

## Introduction

Neurons rely on highly polarised intracellular trafficking to establish and maintain their complex morphology and functional compartmentalisation^1–5^. Directed transport of proteins and organelles from the soma to the axon is essential for axon specification and maintenance^6–8^. Microtubules provide the structural tracks for cargo transport and possess intrinsic polarity that is recognised by molecular motors, enabling directional trafficking within neurons^2,9^. Several regulatory mechanisms have been proposed to govern this process. Notably, axonal trafficking becomes polarised prior to the formation of the axon initial segment (AIS), within a region proximal to the future AIS^7,10^, suggesting that early and spatially restricted mechanisms define axonal transport selectivity.

Increasing evidence indicates that structural microtubule-associated proteins (MAPs) play an active role in polarised trafficking and axon maturation by regulating microtubule organisation and motor-based transport^6,11–13^. Members of the MAP6 and MAP7 families that are spatially restricted to the proximal axon have been shown to regulate axonal cargo entry from the soma and contribute to axon development. For example, MAP6 redistributes from the Golgi and secretory vesicles to the proximal axon as neurons mature^11^, whereas MAP7D2 localises to the proximal axon where it interacts with kinesin-1 to promote axonal trafficking and development^12^. However, whether additional MAPs contribute to soma-to-axon transport through distinct mechanisms, and whether such mechanisms are required to regulate trafficking in mature neurons, remains unclear. In mature hippocampal and sensory neurons, polarised cargo sorting is mediated by a pre-axonal filtering zone located in the proximal axon, characterised by the enrichment of MAP2 and the microtubule bundler TRIM46^6^. Within this region, MAP2 functions as a key regulator of cargo entry by modulating kinesin-1 motor activity, thereby coordinating compartment-specific transport from the soma into the axon^6^. TRIM46, a member of the class I tripartite motif (TRIM) protein family, is an axonal microtubule bundler^14–16^. In developing neurons, TRIM46 exhibits unique microtubule crosslinking activity, promoting the formation of tightly packed, parallel microtubule bundles with a uniform “plus-end-out” orientation^14,17^. This microtubule organisation is thought to provide a directional scaffold for motor-driven cargo transport into the axon^18^. Therefore, TRIM46 plays an important role in organising the proximal axonal microtubule cytoskeleton, supporting axonal transport and the compartmentalisation of axonal and somatodendritic proteins^14,15^.

During neuronal development, TRIM46 is actively transported to the nascent axon via KIF3/KAP3 motor complexes under the control of MARK2 signalling, where it contributes to the establishment of early axonal polarity^19^. However, while this developmental pathway is well characterised, how TRIM46 localisation is maintained in mature axons remains unclear, raising the possibility that distinct regulatory mechanisms operate to preserve axonal organisation beyond development.

MAP1A is one of the most abundantly expressed structural MAPs in the adult mammalian nervous system^20^ and human MAP1A variants have been associated with neurological disease^21,22^. In mice, loss of MAP1A leads to late-onset neuronal death due to impairments in microtubule organisation^23^. However, the cellular mechanisms by which MAP1A maintains neuronal integrity in mature neurons remain poorly understood. Here, we identify a previously unrecognised MAP1A-dependent pathway that operates in mature neurons to maintain TRIM46 localisation in the proximal axon. We show that MAP1A is enriched in the proximal axon and is proteolytically processed by Calpain 10 into heavy and light chains and that this cleavage is required for targeting TRIM46 to this compartment and for maintaining axonal morphology. Mechanistically, the MAP1A light chain interacts with TRIM46 and associates with the tail domain of KIF3 motors, defining a transport complex that facilitates TRIM46 axonal localisation. Together, these results demonstrate that MAP1A processing regulates the molecular organisation of the proximal axon and further supports a model in which distinct MAPs coordinate specific transport routes in mature neurons.

## Method details

### Animals

All experiments with animals were performed in compliance with the guidelines for the welfare of experimental animals issued by the Government of New Zealand and were approved by the Animal Ethical Review Committee of the University of Otago AUP22-44.

### Antibodies and reagents

The following antibodies were used for immunocytochemistry: Rabbit monoclonal anti-MAP1A (Abcam, ab184349), Rabbit monoclonal anti-MAP1A (LC2) (Abcam, ab184350), Mouse monoclonal anti-MAP1A (HC) (Thermo Fisher Scientific, MA5-32988), Mouse monoclonal anti-MAP2 (Sigma-Aldrich, M9942), Guinea pig polyclonal anti-TRIM46 (Synaptic Systems, 377 005), Rabbit polyclonal anti-MAP1B (PA5-78052), Rabbit polyclonal anti-MAP4 (Merck Millipore, AB6020), Rabbit polyclonal anti-MAP6 (Thermo Fisher Scientific, PA5-78399), Rabbit polyclonal anti-MAP7D2 (Sigma-Aldrich, HPA051508), Rabbit polyclonal anti-MAP9 (Bioss, BS-9313R), Mouse monoclonal anti-acetylated tubulin (Sigma-Aldrich, T7451), Rat monoclonal anti-tubulin (tyrosinated) antibody [YL1/2] (Abcam, ab6160), Rabbit polyclonal anti-tubulin β3 (TUBB3) antibody (BioLegend, PRB-435P), Rabbit polyclonal anti-calpain10 (Abcam, ab28226), Rabbit polyclonal anti-KIF3A (Abcam, ab11259), Rabbit polyclonal anti-KIF3B (Thermo Fisher Scientific, PA5-109880), Rabbit polyclonal anti-KIF3C (Thermo Fisher Scientific, PA5-97899), Anti-mouse Alexa488 (Thermo Fisher Scientific, A11029), Anti-rabbit Alexa488 (Thermo Fisher Scientific, A11034), Anti-guinea pig Alexa488 (Thermo Fisher Scientific, A11073), Anti-rat Alexa488 (Thermo Fisher Scientific, A11006), Anti-mouse Alexa568 (Thermo Fisher Scientific, A11031), Anti-rabbit Alexa568 (Thermo Fisher Scientific, A11036), Anti-guinea pig Alexa568 (Thermo Fisher Scientific, A11075), Anti-rat Alexa568 (Thermo Fisher Scientific, A11077), Anti-mouse Alexa647 (Thermo Fisher Scientific, A21236), Anti-rabbit Alexa647 (A21245). The following antibodies were used for Western blot analysis: Rabbit polyclonal anti-GFP (Thermo Fisher Scientific, A-11122), Rabbit polyclonal anti-TagRFP (Evrogen, AB233), Rabbit polyclonal anti-mCherry (Abcam, ab167453), Mouse monoclonal anti-Myc (9E10) (Thermo Fisher Scientific, MA1-980), Donkey anti-mouse IRDye 680RD (LI-COR Biosciences, 926-68072), Donkey anti-rabbit IRDye 800CW (LI-COR Biosciences, 926-32213). Calpeptin was obtained from Sigma-Aldrich (C8999).

### DNA and shRNA constructs

GFP-MAP1A, MAP1A-GFP and GFP-Calpain10 rat sequences were synthesised (e-Zyvec). All other constructs were generated using PCR-based cloning strategies. LC2 and MAP1A-ΔC sequences were PCR-amplified from GFP-MAP1A and subcloned into pEGFP-C1, TagRFP-C1 or mCherry-C2 vectors (Clontech) using primers containing BglII and SalI restriction sites. The Myc epitope sequence (5’GAACAAAAACTCATCTCAGAAGAGGATCTG3’) was generated by annealing complementary oligonucleotides and inserted into pEGFP-LC2 to replace the GFP tag using AgeI and BglII restriction sites. Mouse TRIM46 was PCR-amplified from pcDNA-DEST53-mTrific43 and cloned into pEGFP-C1 or TagRFP-C1 using BglII and SalI restriction sites. TagRFP-Calpain10 was generated by PCR amplification from GFP-Calpain10 followed by subcloning into TagRFP-C1 using BglII and SalI restriction sites. The following shRNA sequences were designed using the Broad Institute Genetic Perturbation platform and Invivogen’s siRNA Wizard online tool: MAP1A shRNA-1 (5’GGCCAGATGAACAAGAAAT3’), MAP1A shRNA-2 (5’ATGTGGATCTTGCCTATAT3’), CAPN10 shRNA (5’GTGCCACACCTCAACTGTT3’), CAPN1 shRNA (5’GAGCAACCCGCAGTTTATT3’), KIF3A shRNA (5’GCCACCATGCCGATCAATA3’), KIF3B shRNA (5’AAAGCTCAAGCATCTTATT3’) and KIF3C shRNA (5’CGAACCGAGCCAAGAACAUUCUCUUGAAAUGUUCUUGGCUCGGUUCGGGG3’).

Complementary oligonucleotides were annealed and ligated into the pSuper vector.

### Sensory neuron culture and electroporation

Sensory neuron cultures were grown as previously described^6^. Briefly, whole dorsal root ganglia were isolated from adult female Sprague Dawley rats (10 weeks or older). Sensory neurons were dissociated with 2mg/ml collagenase type IV (Worthington Biochemical Corp), 1mg/ml trypsin (Sigma-Aldrich) and mechanical trituration. Dissociated neurons were plated on glass coverslips pre-treated with 20 μg/ml poly-D-lysine (Sigma-Aldrich) and 10 μg/ml laminin (Sigma-Aldrich), and neurons were grown in neuron culture media containing DMEM (Thermo Fisher Scientific), 1% FBS (Thermo Fisher Scientific) and 1% penicillin-streptomycin-fungizone (Sigma-Aldrich) at 37°C in 5% CO_2_. Transfection of neurons was performed using the Neon electroporator system (Thermo Fisher Scientific). Neurons were electroporated in suspension, in a 10 μl volume containing ∼1 x10^5^ cells, with 1 μg DNA. For the first 24hrs, transfected neurons were grown in antibiotic-free neuron culture media. Unless otherwise stated, all neurons were fixed at 5DIV using 4% PFA (Sigma-Aldrich). For calpeptin treatment experiments, neurons were treated with 100 μM calpeptin for 48 h prior to fixation with 4% paraformaldehyde (PFA) at 5DIV.

### Primary hippocampal neuron cultures

Hippocampal neuron cultures were grown as previously described^24^. In brief, hippocampi from Sprague Dawley rat pups (0-1 days old) were dissociated with papain (Sigma-Aldrich) and plated at a density of 40,000 cells/coverslip on glass coverslips pre-treated with 20 μg/ml poly-D-lysine (Sigma-Aldrich). Cells were grown in culture media containing Neurobasal A medium (Thermo Fisher Scientific), B27 (Thermo Fisher Scientific), and Glutamax (Thermo Fisher Scientific) and maintained at 37°C in 5% CO_2_ for 21-23DIV prior to fixation using 4% PFA (Sigma-Aldrich).

### HEK293 cell culture and transfection

Human embryonic kidney cells (HEK293, ATCC) were cultured in DMEM/Ham’s F-12 (1:1 ratio, Thermo Fisher Scientific), 10% FBS and 1% penicillin-streptomycin-fungizone at 37°C in 5% CO_2_. HEK293 cells used for immunostainings were first plated on glass coverslips and were transfected using PEI Max (Polysciences) following manufacturer’s instructions. HEK293 cells used for immunoprecipitation lysates were seeded on 100/20 mm cell culture dishes, grown to optimal confluency and transfected using PEI Max.

### Sensory neuron, hippocampal neuron and HEK293 cell immunofluorescence

Neurons and HEK293 cells on glass coverslips were fixed in 4% PFA at room temperature for 15 min. After permeabilising with 0.1% Triton X-100 (Sigma-Aldrich) for 15 min, cells were incubated with primary antibodies in phosphate buffered saline (PBS) and 10% goat serum (Thermo Fisher Scientific) for 1 h, and secondary antibodies in PBS for 1 h. All steps were performed at room temperature, with PBS washes taking place between antibody incubations. Coverslips were mounted on glass slides using Fluorsave (MerckMillipore).

### Whole dorsal root ganglia immunohistochemistry

Whole dorsal root ganglia (DRG) were dissected from adult rats and mice and placed in 4% PFA fixative for 1 h, followed by overnight incubation in 30% sucrose dissolved in phosphate buffered saline (PBS). DRG were embedded in tissue cutting media and frozen at -80° C. Sections were cut at 10 µm using a cryostat and thaw mounted onto Superfrost plus slides (VWR International). Sections were permeabilised and blocked in PBS with 0.3% Trition-X and 10% normal goat serum for 1 h prior to incubation in primary antibodies with 10% normal goat serum and 0.3% Triton-X 100 overnight at 4°C. After washes in PBS sections were incubated in secondary antibodies for 2 h at room temperature. Coverslips were mounted on slides using Fluorosave mounting medium (MerckMillipore).

### Cell/tissue extracts and Immunoblotting

Cell extracts for SDS-PAGE and western blot experiments were prepared from transfected HEK293 cells (either for immunoblotting alone or combined with immunoprecipitation). HEK293 cells transfected with pEGFP (Control), GFP-MAP1A-ΔN and GFP-LC2 (with or without TagRFP-CAPN10 co-transfection) were directly lysed in 2x Laemmli SDS sample buffer, boiled for 5 min at 95°C and loaded into 10% SDS-PAGE gels (Thermo Fisher Scientific). HEK293 cell lysates used for immunoprecipitations were processed in lysis buffer containing 20 mM Tris, 150 mM NaCl, 1% Triton X-100 and protease inhibitor cocktail (Roche). Tissue extracts, or brain lysates, that were used for pulldown-based mass spectrometry were prepared from adult rat brains homogenised in tissue lysis buffer (50 mM Tris-HCl, 150 mM NaCl, 0.1% SDS, 0.2% NP-40, protease inhibitor cocktail).

For western blot assays, samples and PageRuler protein ladder (Thermo Fisher Scientific) were loaded into 10% SDS-PAGE gel, transferred to nitrocellulose membranes and blocked using Intercept TBS blocking buffer (LI-COR Biosciences). Membranes were incubated with primary antibodies overnight at 4°C, and with secondary IRDye antibodies (LI-COR Biosciences) for 1 h at room temperature. Primary and secondary antibodies were made up in blocking buffer, and three washes with TBS with 0.1% Tween-20 were performed after each antibody incubation. Membranes were scanned using an Odyssey Infrared Imaging system and Image Studio Software Version 4 (LI-COR Biosciences).

### GFP and RFP Immunoprecipitation

HEK293 cells processed for immunoprecipitation were first co-transfected with: TagRFP-CAPN10 and pEGFP-C1 (Control) or GFP-MAP1A; GFP-TRIM46 and TagRFP-C1 (Control) or TagRFP-LC2; mCherry-LC2 and pEGFP-C1 (Control) or pEGFP-KIF3(A, B or C)-tail; TagRFP-TRIM46 and pEGFP-C1 (Control) or pEGFP-KIF3(A, B or C)-tail; TagRFP-TRIM46, myc-LC2 and pEGFP-C1 (Control) or pEGFP-KIF3(A, B or C)-tail. HEK293 cells transfected for 24 h were lysed in lysis buffer, centrifuged at 14,000 x g for 15 min at 4°C and incubated with GFP-Trap (Chromotek, gtd-20) or RFP-Trap (Chromotek, rtd-20) magnetic beads following manufacturer’s instructions. Immunoprecipitated proteins retained on the magnetic beads were eluted and processed for SDS-PAGE and western blotting.

### Confocal microscopy of fixed cells and tissue

Images were acquired as z-stacks using an Andor Dragonfly spinning disk confocal system mounted on a Nikon Ti2-E inverted microscope, equipped with 60× (NA 1.49) or 100× (NA 1.45) oil-immersion objectives and an Andor iXon Ultra EMCCD camera. Image acquisition was controlled using Fusion software (version 2.3.0.36; Andor Technology Ltd). Images were processed and analysed using Fiji/ImageJ (NIH) and prepared for presentation using Adobe Illustrator CS6 (Adobe Inc.).

### Live-cell imaging

Live-cell imaging of sensory neurons transfected with mCherry-LC2 was performed on an Andor Dragonfly spinning disk confocal system mounted on a Nikon Ti2-E inverted microscope, equipped with a plan apochromat VC60× (NA 1.2) water-immersion objective and 1.5 magnification and an Andor iXon Ultra EMCCD camera controlled with Fusion software (version 2.3.0.36; Andor Technology Ltd). A 561 nm laser was used for excitation. The microscope was equipped with XY mCherry emission filter. Coverslips with transfected sensory neurons were placed in a microscope holder, fitted into a temperature and CO_2_-regulated stage (kept at 37°C and 5% CO_2_) and supplemented with neuron media. mCherry-LC2 was imaged at 2 second frame rate with 100 millisecond (ms) exposure for 2 minutes.

### Image analysis and quantification

#### Quantification of immunofluorescence

Images were acquired using the same exposure settings and fluorescence intensity was maintained below saturation threshold. Confocal images were processed using maximum intensity projections. For all the analyses, background subtraction of the image was applied. For fluorescence intensity measurements along the axon, line profiles were generated by tracing a segmented line starting at the border point where the soma ends and axon begins and plotting to a distance of 100 μm into the axon. Fluorescence values represented as arbitrary units (A.U). For fluorescence intensity in somas, integrated densities were calculated (intensity/μm^2^, A.U.). For ratio of acetylated-to-tyrosinated tubulin at the proximal axon, intensity measures in a 10 μm^2^ section (5 μm in length) of the proximal axon (closest to the soma) were measured in neurons immunostained for both acetylated and tyrosinated tubulin. All fluorescence values were measured using Fiji/ImageJ (NIH) and averaged over several cells and experiments.

#### Quantification of polarity index

Polarity index was analysed as previously described ^25^. Briefly, average intensities of MAP1A, TRIM46 and Ankyrin-G were analysed in 20 µm sections along axons (at the AIS) and dendrites (two dendrites minimum per cell) of hippocampal neurons (23DIV). Polarity index was determined using the formula (Id-Ia)/(Id+Ia), where Id is mean intensity of dendrite and Ia is mean intensity of axon or AIS. Polarity index values < 0 represent biased polarity towards the AIS while values > 0 biased polarity towards dendrites.

#### Quantification of axon length and width

Montage images of neurons and their respective axons were acquired using Fusion Software (Andor). The segmented line tool of Fiji/Image J software was used to trace and measure the length of the longest neurite per neuron from the axon hillock to the growth cone. Axon width was measured at the start of the proximal axon using GFP as an unbiased fill.

#### Analysis of TRIM46 fragments

Axonal and cell body TRIM46 fragments were categorised into “intact” or “no/fragmented” based on length (μm); fragments >4 μm were classed as intact TRIM46, and fragments <4 μm were identified as no/fragmented TRIM46. When TRIM46 fragments were intact, cells were assigned 100 for “intact” and 0 for “no/fragmented”. When TRIM46 fragments were absent or fragmented, cells were assigned 0 for “intact” and 100 for “no/fragmented”. Scores were analysed as grouped data in GraphPad Prism. Length of TRIM46 fragments in the axon and cell bodies were quantified using Fiji software, while the number of TRIM46 fragments was scored manually.

#### Analysis of LC2 vesicle transport at proximal axon of sensory neurons

Proximal axons of 5DIV sensory neurons expressing mCherry-LC2 were imaged on the Dragonfly spinning disk microscope using 60X water-immersion objective with 1.5x magnification at 2 frame rate for 2mins. Using the segmented line tool in Fiji software, a 100μm-line was traced starting at the beginning of the axon. Lines were used to make kymographs, using Kymoreslicewide plugin (GitHub), which were used to manually quantify the number of non-motile and motile mCherry-LC2 traces (straight and angled traces, respectively) and anterogradely vs retrogradely moving traces (traces moving left-to-right vs right-to-left, respectively).

#### Statistical analysis

Statistical parameters, including the definitions and exact values of n (number of cells), are reported in figures and figure legends. Data and statistical analysis were performed with Excel (Microsoft) and GraphPad Prism software (GraphPad Software INC). The assumption of data normality was checked using D’Agostino-Pearson omnibus test. Statistical analyses used were: Ordinary one-way ANOVA with Dunnett’s multiple comparisons post-hoc analysis for normal distributions or Kruskal-Wallis test with post-hoc Dunn’s multiple comparisons test for non-normal distributions (not significant is p > 0.05, *p < 0.05, **p < 0.01, ***p < 0.001). Graphs were made using GraphPad Prism software.

## Results

### MAP1A is specifically enriched in the proximal axon of mature neurons

Spatially restricted structural MAPs regulate trafficking and axon maturation in developing neurons^6,11–13^. However, the localisation and function of these MAPs in mature neurons remain poorly defined. To address this, we selected a panel of structural MAPs representing major MAP families with established roles in neuronal microtubule organisation and trafficking, including MAP1A^26,27^, MAP1B^27,28^, MAP2^6^, MAP4^29^, MAP6^11^, MAP7D2^12^ and MAP9^30^, and examined their endogenous distribution by immunofluorescence in cultured rat adult sensory neurons, alongside TRIM46 as a marker of the proximal axon^6,14^ (Figure 1A). Quantitative fluorescence intensity analysis showed that MAP1A, MAP2, and TRIM46 were enriched at the proximal axon, whereas the other MAPs examined were uniformly distributed (Figure 1B). Line profile analysis showed that MAP1A fluorescence intensity was centred within the MAP2- and TRIM46-enriched region and overlapped with both proteins (Figures 1C). To determine whether this localisation pattern is conserved across neuronal types, we next examined mature central nervous system neurons. Consistent with previous reports ^27,31^, polarity index analysis revealed strong dendritic localisation of MAP1A in mature hippocampal neurons (Figure 1D). However, MAP1A was also enriched in the proximal axon, close to the AIS marker Ankyrin G, and partially overlapping with TRIM46 (Figure 1E), in line with our observations in sensory neurons. To gain more insight into how proximal axon organisation is spatially and temporally coordinated, we performed a time course analysis of endogenous MAP1A and TRIM46 distribution during sensory axon outgrowth (Figure 1F). Quantitative analysis revealed that at 1 day in vitro (1 DIV), neither MAP1A nor TRIM46 was localised to the proximal axon. By 2 DIV, MAP1A was enriched in the axon, whereas TRIM46 remained absent, indicating that MAP1A enrichment precedes TRIM46 localisation to this region (Figure 1F). At 3–4 DIV, MAP1A remained enriched in the proximal axon, and discrete TRIM46 fragments began to appear (Figure 1F). By 5 DIV, both MAP1A and TRIM46 showed clear enrichment and compartmentalisation within the proximal axon (Figure 1F). In agreement with these observations in cultured sensory neurons, MAP1A was also enriched in the proximal axon of adult sensory neurons in vivo, where it partially overlapped with TRIM46 (Figure 1G). Together, these findings identify MAP1A as an early and proximally enriched MAP in mature axons.

**Figure 1.**
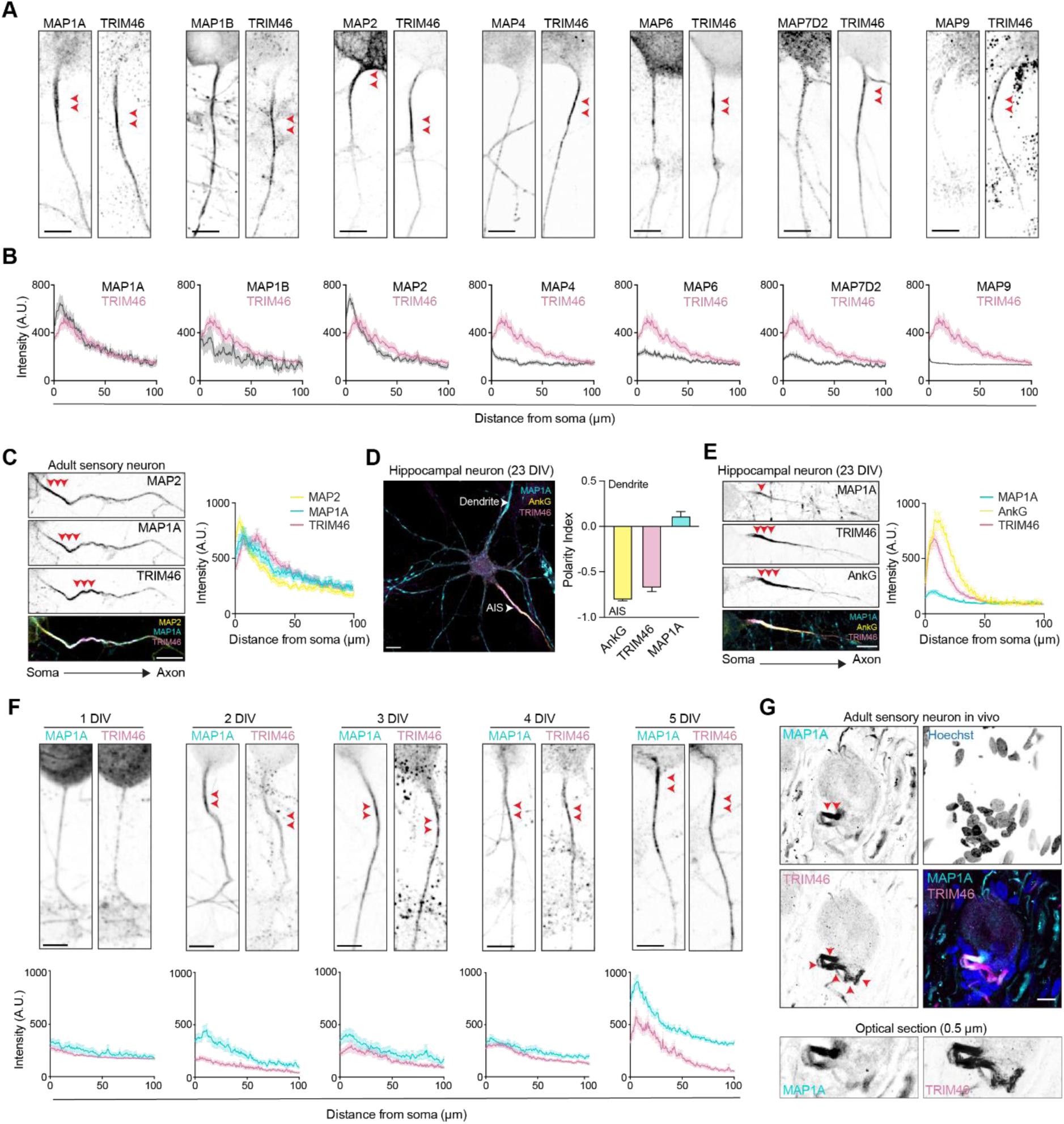
MAP1A localises at the proximal axon of mature neurons. (A) Representative images of adult sensory neurons at 5 days in vitro (DIV) immunostained with the indicated antibodies. Red arrowheads indicate the specific staining. (B) Average fluorescent intensity of MAP1A (n=78), MAP1B (n=23), MAP2 (n=78), MAP4 (n=22), MAP6 (n=29), MAP72D (n=23), MAP9 (n=16) and TRIM46 (n=52) in the proximal axon of 5 DIV adult sensory neurons. (C) Representative adult sensory neuron (5 DIV) immunostained with the indicated antibodies. Red arrowheads indicate the specific staining in the proximal axon. The graph shows the average fluorescent intensity of MAP2 (n=78), MAP1A (n=78) and TRIM46 (n=78) along the proximal axon of 5DIV adult sensory neurons. (D) Representative mature hippocampal neuron immunostained with the indicated antibodies. White arrowheads indicate the dendrites and the axon initial segment (AIS). Graph shows the mean ± SEM polarity index of Ankyrin G (n=50), TRIM46 (n=50) and MAP1A (n=50). (E) Representative image of the AIS of a mature hippocampal neuron immunostained with the indicated antibodies. The graph shows the average fluorescent intensity of MAP1A (n=53), Ankyrin G (n=53) and TRIM46 (n=53) along the proximal axon. (F) Representative images of adult sensory neurons immunostained with the indicated antibodies at the specified time points. Graphs show the proximal axon average fluorescent intensities at 1 DIV (MAP1A n=62, TRIM46 n=71), 2 DIV (MAP1A n=77, TRIM46 n=59), 3 DIV (MAP1A n=58, TRIM46 n=58), 4 DIV (MAP1A n=88, TRIM46 n=72), 5 DIV (MAP1A n=87, TRIM46 n=58. (G) Immunohistochemistry of MAP1A and TRIM46 in adult sensory neurons in vivo. n= neurons, A.U., arbitrary units. Mean ± SEM; at least three independent experiments. Scale bars indicate 10 µm in (A), (C), (D), (E) and (G).

### MAP1A is required for the proper localisation of TRIM46 at the proximal axon

To investigate the role of MAP1A in mature neurons, we silenced MAP1A expression in adult sensory neurons using a short hairpin RNA (shRNA) knockdown approach. Knockdown efficiency of endogenous MAP1A using two independent shRNAs targeting either the N-terminal or C-terminal coding regions of the MAP1A transcript was validated by immunocytochemistry. MAP1A immunofluorescence was significantly reduced in both the soma and proximal axon following treatment with either shRNA (Figure S1A-E). MAP1A depletion significantly reduced axon length (Figure 2A, B) and caused an abnormal enlargement of the proximal axon width (Figure 2C, D), consistent with in vivo observations^23^. We hypothesised that these axonal enlargements in MAP1A-depleted neurons may reflect underlying defects in cytoskeletal organisation. Indeed, previous studies showed that homozygous MAP1A mutant mice exhibit abnormal dendritic shaft swellings and structural abnormalities in the proximal axon^23^. Furthermore, disruption of microtubule organisation, either through treatment with depolymerising agents such as nocodazole^32^ or by silencing the dynein regulator NDEL1^33^, also leads to axonal enlargement. Given that MAP1A localises to the proximal axon and partially overlaps with TRIM46 (Figure 1F, G), and TRIM46 is a key microtubule-bundling protein^14^, we asked whether loss of MAP1A alters TRIM46 localisation and contributes to proximal axon enlargement. We therefore analysed the localisation and distribution of TRIM46 in the soma and axon of adult sensory neurons following MAP1A knockdown (Figure 2E). In control neurons, TRIM46 was excluded from the soma and formed a single continuous enrichment in the proximal axon (Figure 2E–L). Following MAP1A depletion, this organisation was disrupted, with increased TRIM46-positive fragments in the soma and a fragmented or absent proximal axon TRIM46 signal (Figure 2E-L). These findings indicate that MAP1A is required for proper TRIM46 localisation to the proximal axon, and that its loss leads to TRIM46 mislocalisation to the soma, potentially contributing to microtubule disorganisation and axonal enlargement.

**Figure 2.**
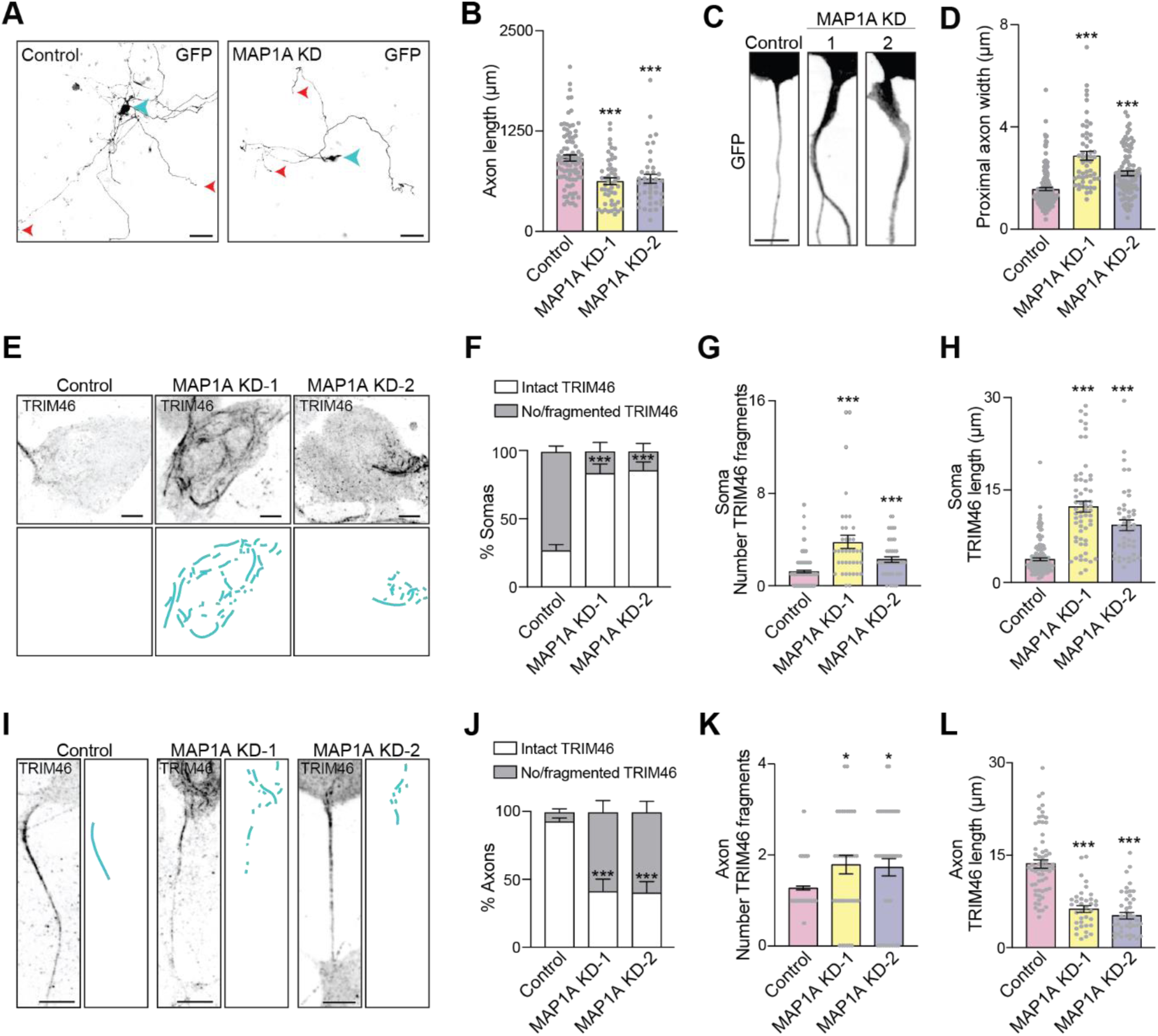
MAP1A regulates the localisation of TRIM46 at the proximal axon of mature neurons. (A) Representative images of control and MAP1A knockdown (KD) adult sensory neurons (5DIV). Blue arrowheads indicate the soma and red arrowheads indicate the axon tips. (B) Quantification of axon length under the indicated conditions (control n=88, MAP1A KD-1 n=49, MAP1A KD-2 n=40). (C) Representative images of the proximal axon of control and MAP1A KD adult sensory neurons (5DIV). (D) Quantification of proximal axon width (control n=171, MAP1A KD-1 n=53, MAP1A KD-2 n=95). (E) Representative images of the soma of adult sensory neurons (5DIV) immunostained for TRIM46 under the indicated conditions. Panels with blue traces show TRIM46 distribution in the soma. (F) Percentage of somata with intact TRIM46 stretches (> 4 µm fragment length) or absent/fragmented (< 4 µm fragment length) under the indicated conditions (control n=140, MAP1A KD-1 n=38, MAP1A KD-2 n=44). (G) Mean number of TRIM46 fragments in the soma (control n=166, MAP1A KD-1 n=39, MAP1A KD-2 n=44). (H) Mean length of TRIM46 fragments in the soma (control n=164, MAP1A KD-1 n=59, MAP1A KD-2 n=44). (I) Representative images of the proximal of adult sensory neurons (5DIV) immunostained for TRIM46 under the indicated conditions. Panels with blue traces show TRIM46 distribution in the proximal axon. (J) Percentage of axons with intact TRIM46 stretches (> 4 µm fragment length) or absent/fragmented (< 4 µm fragment length) under the indicated conditions (control n=163, MAP1A KD-1 n=38, MAP1A KD-2 n=44). (K) Mean number of TRIM46 fragments in the proximal axon (control n=163, MAP1A KD-1 n=38, MAP1A KD-2 n=44). (L) Mean length of TRIM46 fragments in the proximal axon (control n=63, MAP1A KD-1 n=36, MAP1A KD-2 n=44). n= neurons, A.U., arbitrary units. Data displayed as mean ± SEM. Scale bars indicate 100 µm in (A) and 10 µm in (E) and (I). ***p <0.001; **p < 0.01; *p <0.05 (B), (D), (F)-(H) and (J)-(L) Kruskal-Wallis test, with Dunn’s multiple comparisons post-hoc analysis.

### TRIM46 localisation at the proximal axon depends on its interaction with MAP1A light chain 2

The *MAP1A* gene encodes a precursor polypeptide that is proteolytically cleaved to produce a MAP1A heavy chain (HC) and a light chain (LC2)^34^. To further elucidate the mechanism by which MAP1A regulates TRIM46 localisation, we performed rescue experiments to identify the domains of MAP1A that are involved. Two shRNAs targeting distinct regions of the MAP1A transcript were used: one directed against the HC-coding sequence and the other against the LC2-coding sequence (Figure S1A). This strategy enabled depletion of endogenous full-length MAP1A while permitting expression of GFP-tagged rescue constructs corresponding to the non-targeted domain. Adult sensory neurons were transfected with either HC-targeting shRNA together with GFP-LC2, or LC2-targeting shRNA together with GFP-HC, followed by immunostaining for endogenous TRIM46 (Figure 3A). As previously observed, MAP1A knockdown increased the number of TRIM46-positive fragments in the soma compared to control neurons; however, this phenotype was rescued by LC2, but not by HC expression (Figure 3A, B). Line profile analysis further showed that MAP1A depletion impaired TRIM46 enrichment in the proximal axon, whereas expression of GFP-LC2 was sufficient to restore TRIM46 levels to those of controls (Figure 3C, D). Consistently, MAP1A silencing significantly reduced the proportion of axons displaying intact TRIM46 (Figure 3E) and increased the number of TRIM46-positive fragments in the axon (Figure 3F); both defects were rescued by GFP-LC2, but not GFP-HC (Figure 3E, F).

**Figure 3.**
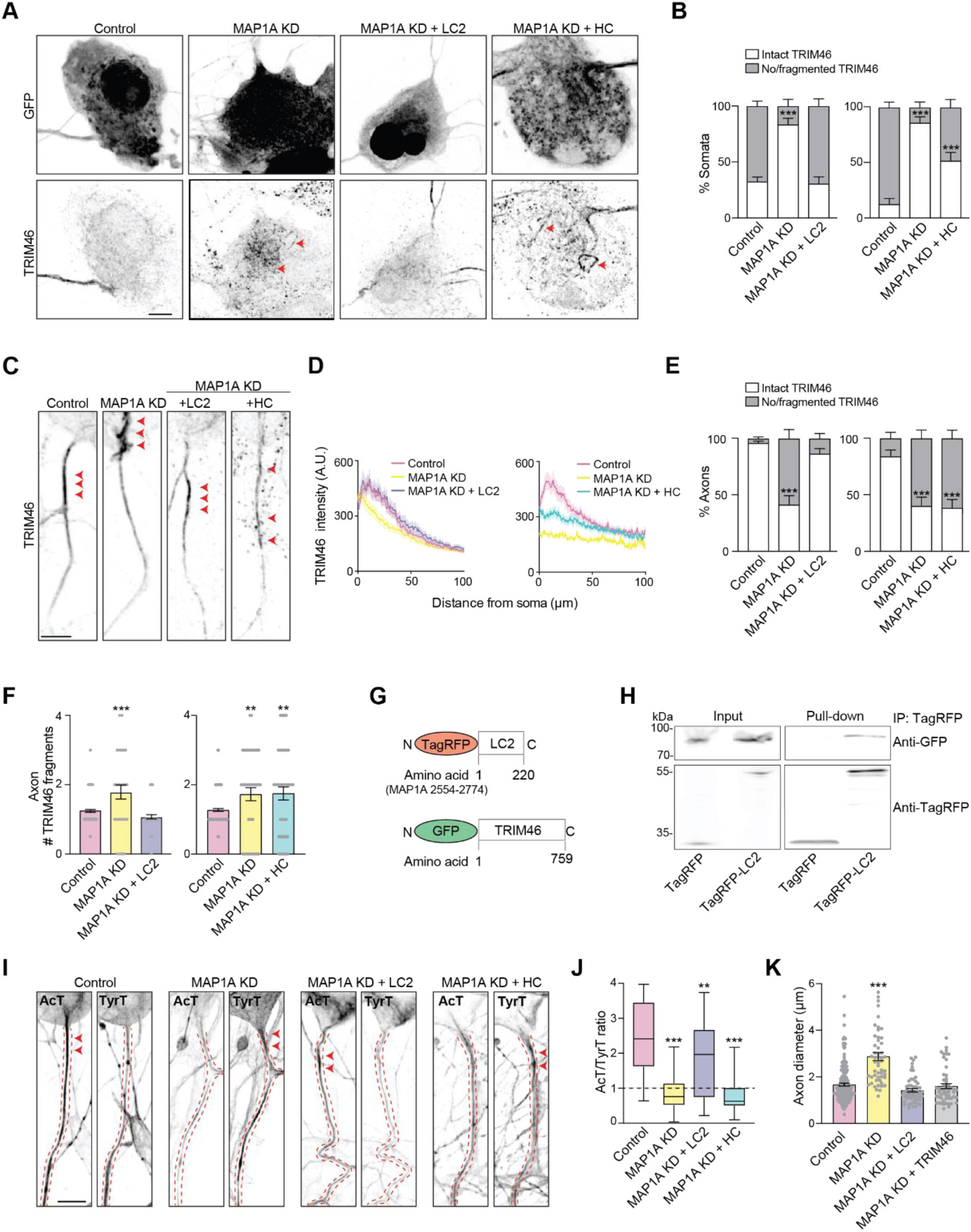
MAP1A light chain 2 interacts with TRIM46 to regulate its localisation and maintain axon morphology. (A) Representative images of adult sensory neuron somas (5 DIV) under the indicated conditions and immunostained for endogenous TRIM46. Red arrowheads indicate TRIM46 fragments in the soma. (B) Percentage of somata with intact TRIM46 stretches (> 4 µm fragment length) or absent/fragmented (< 4 µm fragment length) under the indicated conditions (left graph, control n=140, MAP1A KD n=38, MAP1A KD + LC2 n=54; right graph, control n=140, MAP1A KD n=44, MAP1A KD + HC n=46). (C) Representative images of adult sensory neuron proximal axons (5 DIV) under the indicated conditions and immunostained for endogenous TRIM46. Red arrowheads indicate TRIM46 fragments in the axon. (D) Mean fluorescence intensity of TRIM46 along the proximal axon under the indicated conditions (left graph, control n=126, MAP1A KD n=131, MAP1A KD + LC2 n=94; right graph, control n=132, MAP1A KD n=32, MAP1A KD + HC n=92). (E) Percentage of axons with intact TRIM46 stretches (> 4 µm fragment length) or absent/fragmented (< 4 µm fragment length) under the indicated conditions (left graph, control n=163, MAP1A KD n=38, MAP1A KD + LC2 n=54; right graph, control n=163, MAP1A KD n=44, MAP1A KD + HC n=46). (F) Mean number of TRIM46 fragments in the proximal axon (left graph, control n=163, MAP1A KD n=38, MAP1A KD + LC2 n=54; right graph, control n=163, MAP1A KD n=44, MAP1A KD + HC n=46). (G) Schematics of the indicated constructs. (H) Western blot with indicated antibodies of TagRFP-LC2 immunoprecipitations. (I) Representative images of adult sensory neurons (5 DIV) immunostained for acetylated tubulin (AcT) and tyrosinated tubulin (TyrT) under the indicated conditions. Dashed red lines outline the proximal axon. (J) Average ratio the acetylated to tyrosinated tubulin at the proximal axon under the indicated conditions (control n=43, MAP1A KD n=46, MAP1A KD + LC2 n=40, MAP1A KD + HC n=40). (K) Quantification of proximal axon width under the indicated conditions (control n=167, MAP1A KD n=53, MAP1A KD + LC2 n=53, MAP1A KD + TRIM46 n=54). n=neurons, A.U., arbitrary units. Mean ± SEM; at least three independent experiments. Scale bars indicate 10 µm in (A), (C) and (I). ***p <0.001; **p < 0.01; *p <0.05 (B), (E), (I) and (J) Kruskal-Wallis test, with Dunn’s multiple comparisons post-hoc analysis; (F) Ordinary one-way ANOVA with Dunnett’s multiple comparisons post-hoc analysis.

To determine whether this rescue reflects a direct association between LC2 and TRIM46, we performed biochemical interaction assays using co-immunoprecipitation from HEK293 cells expressing both proteins (Figure 3G, H). These experiments demonstrated that LC2 robustly co-precipitates TRIM46, indicating an interaction between the two proteins (Figure 3H). Given the role of TRIM46 in microtubule bundling and stabilisation^14,16,17^, we next investigated whether MAP1A depletion alters microtubule stability in the proximal axon. Immunostaining of control neurons, MAP1A knockdown neurons, and MAP1A knockdown neurons rescued with either GFP-HC or GFP-LC2 was performed using antibodies against acetylated and tyrosinated tubulin as markers of stable and dynamic microtubule populations, respectively (Figure 3I). Quantification of the ratio of acetylated to tyrosinated tubulin revealed that MAP1A knockdown significantly reduced microtubule stability (Figure 3I, J). In line with the TRIM46 axonal localisation data (Figure 3C-F), GFP-LC2 expression restored acetylated tubulin levels to that of control neurons, indicating recovery of microtubule stability (Figure 3I, J). Given that MAP1A depletion disrupts TRIM46 localisation in the proximal axon and reduces microtubule stability, we next tested whether restoring LC2 function or TRIM46 localisation was sufficient to rescue the associated axonal morphological phenotype. As previously observed, MAP1A knockdown resulted in a significant increase in proximal axon width (Figure 2C, D and Figure 3K). However, expression of either GFP-LC2 or GFP-TRIM46 restored axon thickness to levels comparable to control neurons (Figure 3K). Together, these results indicate that MAP1A LC2 is both necessary and sufficient to restore TRIM46 localisation, through a direct interaction with TRIM46, and to maintain microtubule organisation and proximal axon morphology.

### Cleavage of MAP1A by Calpain-10 is necessary for TRIM46 localisation to the proximal axon

MAP1A family proteins have recently been identified as substrates of Calpain-10 (CAPN10)^35^. Although other Calpains have been implicated in neuronal development^36^, the role of CAPN10 in neurons is largely unknown. To determine whether proteolytic processing of MAP1A is required for TRIM46 localisation in the proximal axon, we first examined the distribution of the HC and LC2 domains in mature sensory neurons using domain-specific antibodies. While HC was enriched in the proximal axon (Figure 4A), as determined by line-profile analysis of fluorescence intensity (Figure 4B), LC2 showed a punctate distribution throughout the soma (Figure 4A, B). This differential localisation suggests that MAP1A undergoes spatially regulated proteolytic processing in neurons. Notably, CAPN10 was also enriched in the proximal axon (Figure 4C) and its fluorescence intensity profile overlapped with that of HC (Figure 4D). To test the function of CAPN10, we inhibited calpain activity using the broad-spectrum calpain inhibitor calpeptin or specifically silenced CAPN10 using an shRNA (Figure 4E). Given that both manipulations impaired MAP1A processing, we next examined whether TRIM46 localisation was affected. Critically, both calpeptin treatment and CAPN10 knockdown, but not silencing of the related calpain CAPN1, prevented TRIM46 enrichment in the proximal axon (Figure 4F) as shown by the line profiles (Figure 4G). Together, these results indicate that specific CAPN10-mediated cleavage of MAP1A is required to enable TRIM46 localisation to the proximal axon.

**Figure 4.**
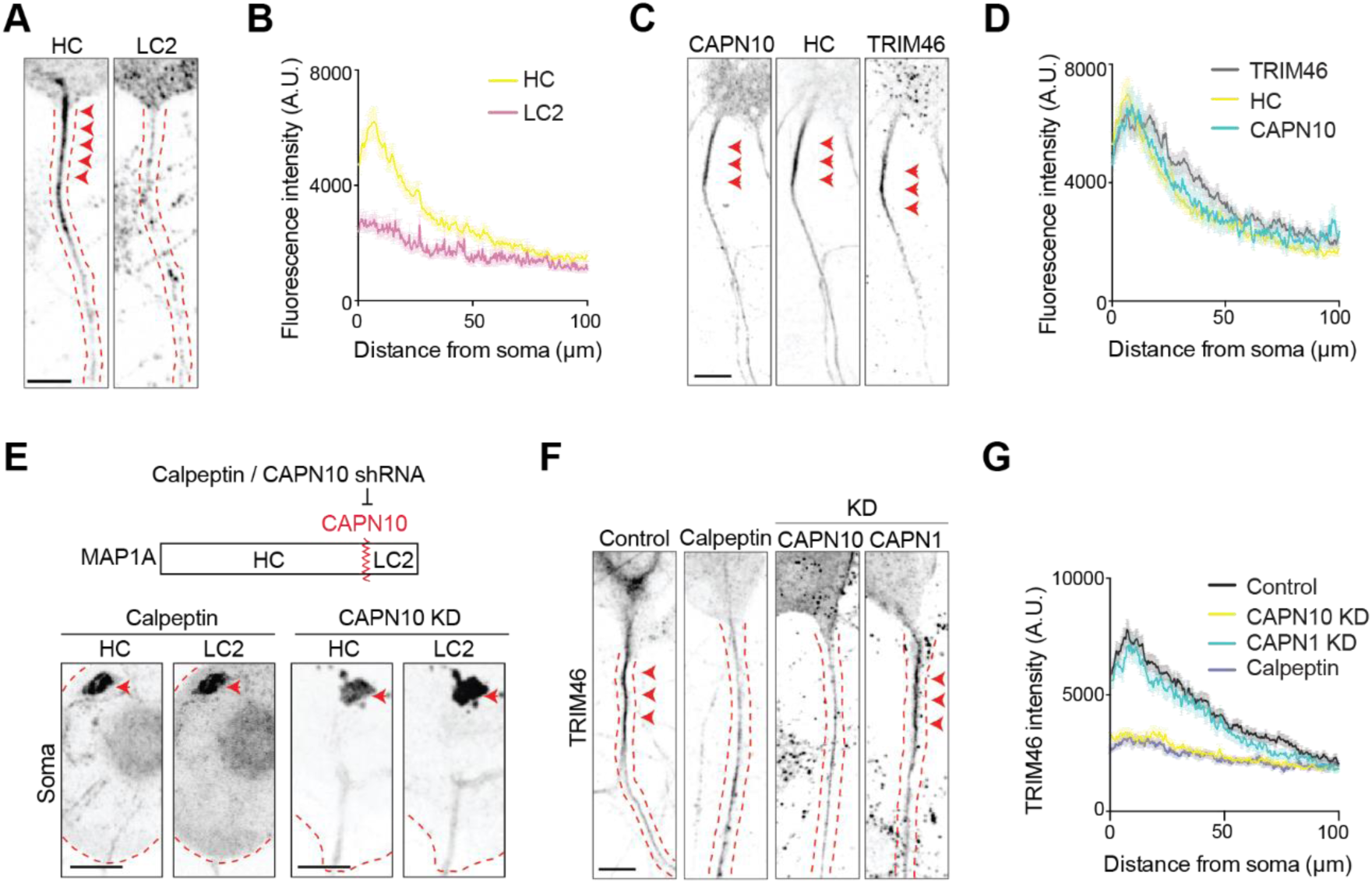
Calpain-10–mediated cleavage of MAP1A is required for TRIM46 localisation to the proximal axon. (A) Representative images of adult sensory proximal axons (5 DIV) immunostained with specific antibodies recognising the HC or LC2 domains of MAP2. Red arrowheads indicate protein distribution in the proximal axon. (B) Average fluorescent intensity of HC or LC2 along the proximal axon (n=66). (C) Representative images of the proximal axon of adult sensory neurons (5 DIV) immunostained with the indicated antibodies. Red arrowheads indicate protein distribution in the proximal axon. (D) Average fluorescent intensity of endogenous Calpain-10 (CAPN10), HC or TRIM46 along the proximal axon (n=116). (E) Top, Schematic depicting inhibition of MAP1A cleavage by calpeptin or CAPN10 knockdown. Bottom, representative images of somata from adult sensory neurons at 5 DIV under the indicated conditions (KD = knockdown), immunostained for endogenous HC and LC2. Red arrowheads denote sites of HC and LC2 aggregation and co-localisation. (F) Representative images of the proximal axon of adult sensory neurons (5 DIV) immunostained for endogenous TRIM46 under the indicated conditions (KD=knockdown). Red arrowheads indicate enriched TRIM46 in the proximal axon. (G) Average fluorescent intensity of endogenous TRIM46 along the proximal axon under the indicated conditions (n=105). n= neurons, A.U., arbitrary units. Mean ± SEM; at least three independent experiments. Scale bars indicate 10 µm in (A), (C), (E) and (F).

### LC2 interacts with kinesin-2 motors to transport TRIM46 to the proximal axon

Having established that MAP1A cleavage is required for LC2-dependent localisation of TRIM46 to the proximal axon, we next sought to determine how LC2 mediates this process. Given the importance of motor-driven transport in the delivery of axonal proteins, we tested whether LC2 couples TRIM46 to microtubule-based transport machinery. Previous work has shown that kinesin-2 family members (KIF3) regulate TRIM46 transport in developing neurons via MARK2 signalling^19^. Based on this, we examined whether KIF3 motors are required for TRIM46 localisation in mature neurons by a knockdown approach. The efficiency of KIF3 motor knockdown was confirmed by immunocytochemistry (Figure S2). Line profile analysis showed that depletion of KIF3A and KIF3C, but not KIF3B, significantly disrupted TRIM46 enrichment in the proximal axon (Figure 5A, B) and reduced the proportion of axons with intact TRIM46. (Figure 5C), indicating that kinesin-2 motors are also required for TRIM46 transport in mature neurons. We next investigated whether KIF3 motors are required for LC2 trafficking to the proximal axon. Live-cell imaging and kymograph analysis revealed an increase in static, non-motile mCherry-LC2 vesicles in KIF3A- or KIF3C-depleted neurons, but not in KIF3B-depleted neurons, compared to controls (Figure 5D, E). This was accompanied by a reduction in anterograde vesicle transport from the soma into the axon (Figure 5F), indicating impaired LC2 transport. Given that LC2 interacts with TRIM46 (Figure 3H) and that kinesin-2 motors are required for both LC2 trafficking and TRIM46 localisation, we next investigated whether LC2 links TRIM46 to kinesin-2 motors. As HEK293 cells do not express endogenous MAP1A, they provide a useful system to determine whether LC2 is sufficient to mediate interactions between TRIM46 and KIF3 motors. Pull-down experiments revealed that the tail domains of all three KIF3 motors co-precipitated LC2 (Figure 5G, H). In contrast, KIF3 tail constructs failed to co-precipitate TRIM46 when expressed in the absence of LC2 (Figure 5I), suggesting that TRIM46 does not interact directly with kinesin-2 motors. Therefore, we hypothesised that LC2 mediates the association between TRIM46 and kinesin-2. Consistent with this model, co-expression of GFP-KIF3 tail constructs, TagRFP-TRIM46, and Myc-LC2 resulted in co-precipitation of both TRIM46 and LC2 with each kinesin-2 motor tail (Figure 6J), indicating that LC2 is sufficient to link TRIM46 to kinesin-2 motors. Together, these results demonstrate that in mature neurons TRIM46 associates with KIF3A/C motors via LC2, which acts as an adaptor coupling TRIM46 to kinesin-driven transport. Overall, our data support a model in which CAPN10-mediated cleavage of MAP1A generates LC2, enabling the formation of a TRIM46–LC2 complex that is transported to the proximal axon by the KIF3A/KIF3C heterodimer motor (Figure 5K).

**Figure 5.**
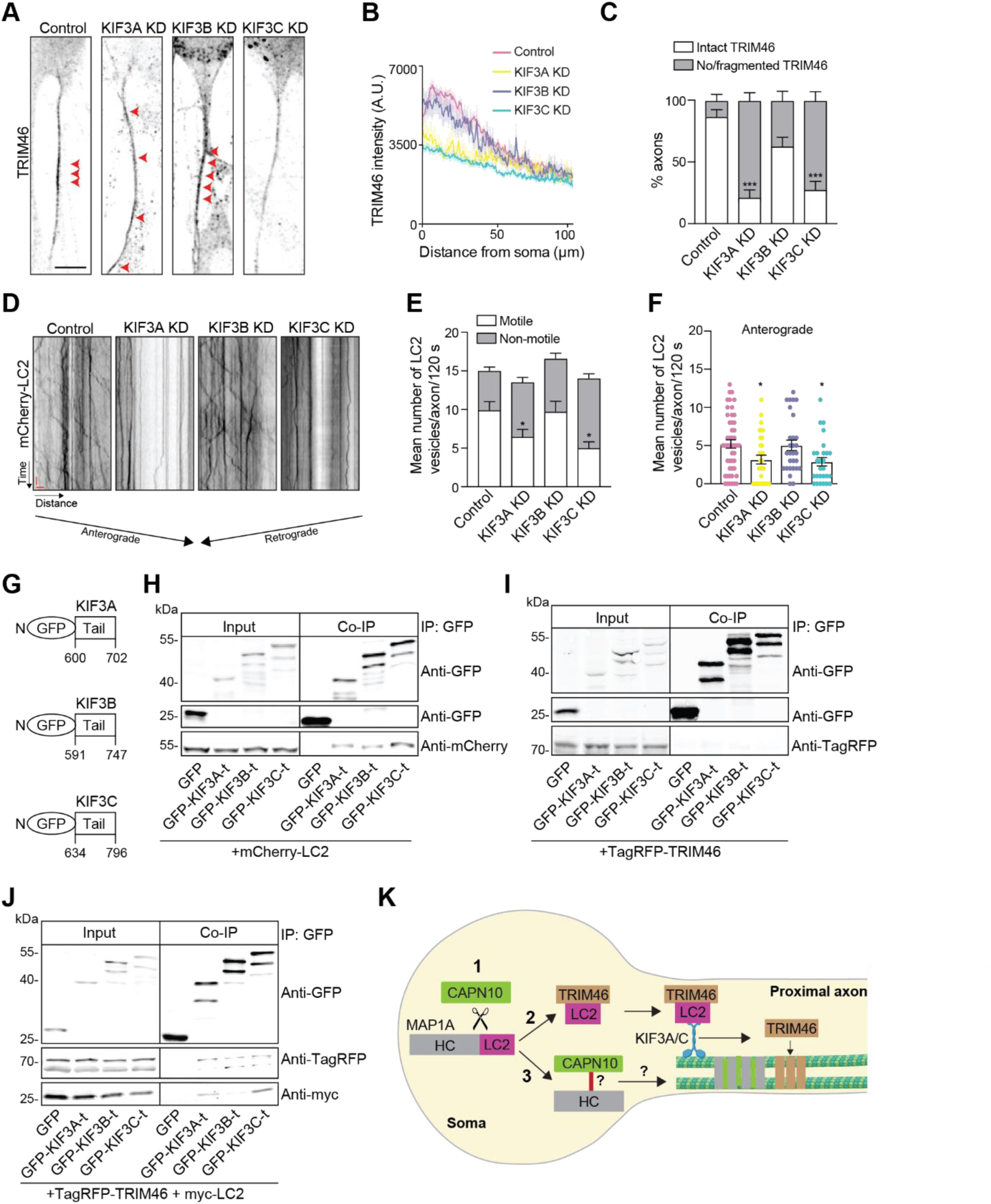
LC2 couples TRIM46 to kinesin-2 motors for delivery to the proximal axon. (A) Representative images of the proximal axon of adult sensory neurons (5 DIV) immunostained for endogenous TRIM46 under the indicated conditions (KD= knockdown). Red arrowheads indicate TRIM46 localisation. (B) Average fluorescent intensity of endogenous TRIM46 along the proximal axon under the indicated conditions (control n=119, KIF3A KD n=119, KIF3B KD n=107, KIF3C KD n=169). (C) Percentage of axons with intact TRIM46 stretches (> 4 µm fragment length) or absent/fragmented (< 4 µm fragment length) under the indicated conditions (control n=39, KIF3A KD n=46, KIF3B KD n=46, KIF3C KD n=43). (D) Representative kymographs of mCherry-LC2 vesicles in the proximal axon under the indicated conditions. Scale bars time y-axis 10 s and distance x-axis 5 µm. (E) Average number of motile and non-motile mCherry-LC2 vesicles in the proximal axon (100 µm) under the indicated conditions (control n= 48, KIF3A KD n=31, KIF3B KD n=29, KIF3C KD n=27). (F) Mean number of anterogradely moving mCherry-LC2 in the proximal axon (control n= 48, KIF3A KD n=31, KIF3B KD n=29, KIF3C KD n=27). (G) Schematic of the indicated constructs. (H-J) Western blot analysis of GFP–KIF3A/B/C tail (GFP–KIF3A/B/C-t) co-immunoprecipitates under the indicated conditions, probed with the indicated antibodies. (K) Graphical model showing (1) MAP1A cleavage by calpain-10 (CAPN10), releasing LC2; (2) LC2 interaction with TRIM46 and kinesin-2 motors to mediate TRIM46 transport to the proximal axon and (3) transport of CAPN10 and the MAP1A heavy chain (HC) to the proximal axon by an unknown mechanism. n= neurons, A.U., arbitrary units. Mean ± SEM; at least three independent experiments. Scale bars indicate 10 µm in (A). ***p <0.001; **p < 0.01; *p <0.05 (C) Kruskal-Wallis test, with Dunn’s multiple comparisons post-hoc analysis; (E) and (F) Ordinary one-way ANOVA with Dunnett’s multiple comparisons post-hoc analysis.

## Discussion

The maintenance of neuronal polarity in mature neurons requires continuous coordination between intracellular transport and cytoskeletal organisation, yet the mechanisms that sustain protein localisation at the proximal axon remain poorly understood. Here, we identify a proteolysis-dependent pathway that links these processes, in which Calpain-10-mediated cleavage of MAP1A generates LC2, a transport adaptor that couples TRIM46 to kinesin-2 motors to ensure its localisation to the proximal axon and thereby maintain axonal organisation. In contrast to mechanisms described during development, our findings define a pathway that specifically operates to preserve axonal architecture in mature neurons.

MAP1A is a member of the MAP1 family, which also includes MAP1B, and is highly expressed in the dendrites of mature neurons^34^. It has been implicated in activity-dependent dendritic branching and arbor stabilisation^31^, and in vivo studies have established its role in organising the somatodendritic microtubule network^23^. Our findings extend this view by showing that MAP1A is also enriched at the proximal axon in mature hippocampal neurons and in adult sensory neurons, which lack a classical dendritic compartment. This localisation is consistent with previous proximity labelling studies in DIV14 hippocampal neurons, which identified MAP1A in the axon initial segment in close proximity to NDEL^37^. Similarly, MAP1A enrichment at the proximal axon has been reported in Purkinje cells, where it becomes apparent following neuronal maturation^23^. Together, these observations indicate that MAP1A localisation at the mature proximal axon is conserved across multiple neuronal subtypes and are consistent with a model in which MAP1A contributes to the maintenance of the axon rather than its initial specification and development. In line with this idea, MAP1A depletion in adult sensory neurons resulted in abnormal proximal axon morphology, characterised by increased axonal width. Comparable phenotypes have been reported in MAP1A homozygous mutant mice, which carry a deletion of 297 amino acids from the C-terminus of the MAP1A heavy chain together with loss of the entire LC2 light chain. In these animals, axonal swellings emerge after neuronal maturation and are associated with disrupted microtubule organisation^23^. These findings align with previous work showing that selective destabilisation of microtubules at the proximal axon is sufficient to induce axonal enlargement^32,33^, highlighting the sensitivity of this compartment to perturbations in cytoskeletal organisation. Together, these results position MAP1A as a key regulator of microtubule integrity required for maintaining proximal axon structure.

A key finding of this study is the identification of a proteolysis-dependent mechanism underlying MAP1A function in mature neurons. While MAP1 family proteins are known to undergo cleavage^20,34,35^, the functional significance of this processing has remained unclear. More broadly, calpains have been implicated in neuronal morphological remodelling, yet the roles of individual family members remain poorly defined^36^. Our findings identify Calpain-10 as a previously unrecognised regulator in neurons and suggest that selective proteolysis of MAPs can actively control protein localisation rather than simply mediate protein turnover^30^.

In this context, MAP1A cleavage generates LC2, which functions as a transport adaptor linking TRIM46 to kinesin-2 motors. TRIM46 is a key organiser of parallel microtubule bundles at the proximal axon and is important for axonal transport and the compartmentalisation of axonal and somatodendritic proteins^14,16,17^. In developing neurons efficient delivery of TRIM46 to the proximal axon depends on MARK2 signalling^19^. Our findings expand this model by identifying a distinct, proteolysis-dependent mechanism that maintains TRIM46 localisation in mature neurons, thereby linking MAP1A processing to microtubule organisation and the maintenance of axonal morphology.

LC2 has previously been implicated in the trafficking of diverse cargoes, including PSD-93/95, RhoB, ion channels, and AMPA receptor-associated proteins^26,38–42^, suggesting a general role in coordinating protein localisation. However, the motor proteins underlying LC2-dependent transport have not been defined. Our results address this gap by identifying kinesin-2 motors, specifically KIF3A and KIF3C, as key mediators of LC2 trafficking in mature neurons. In our system, overexpression of LC2 was sufficient to rescue the axonal morphology defects induced by MAP1A depletion, indicating that LC2-mediated transport of MAPs is a key determinant of proximal axon organisation. In contrast, MAP1A mutant mice exhibit axonal swellings and progressive degeneration of Purkinje cells that is not rescued by LC2^23^, suggesting that full-length MAP1A or the heavy chain fulfil additional functions beyond those mediated by the light chain.

We further found that both the MAP1A heavy chain and calpain-10 also localise to the proximal axon following injury. However, the mechanism by which they are recruited, their functional roles at this site and whether they interact with LC2 or with each other remain unknown. One possibility is that calpain-10 is transported as part of a MAP1A-associated complex, or that after MAP1A cleavage in the soma, calpain-10 remains bound to the heavy chain fragment and are selectively trafficked into the axon. Together, these observations suggest that MAP1A operates through functionally distinct domains that differentially regulate axonal organisation and neuronal survival in a context-dependent manner. One possible explanation is that LC2 primarily supports cargo transport at the proximal axon, whereas full-length MAP1A, or its heavy chain, may be required for maintaining the integrity of the somatodendritic cytoskeleton and overall neuronal viability. These distinctions may be particularly relevant in Purkinje cells, which possess elaborate dendritic arbours. Future work will be necessary to define the role of the MAP1A heavy chain and calpain-10 at the proximal axon and to determine whether they function within a shared pathway.

The functional relevance of the MAP1A pathway is underscored by its potential link to human disease. Truncating mutations in MAP1A have been associated with neurodevelopmental disorders, including autism spectrum disorder and attention-deficit/hyperactivity disorder^21,22^. Given that such mutations are likely to disrupt MAP1A processing, it is plausible that impaired generation of LC2 and consequent defects in TRIM46 transport contribute to disease pathology. Notably, sensory abnormalities are a prominent feature of autism spectrum disorder^43^, raising the possibility that disruption of MAP1A-dependent transport in sensory neurons contributes to altered sensory function^44,45^.

More broadly, our findings suggest that proteolytic processing of cytoskeletal proteins represents a regulatory mechanism for controlling intracellular transport. Microtubule-associated proteins have traditionally been viewed as structural regulators; however, increasing evidence indicates that they also participate in coordinating motor activity and in defining polarized cargo trafficking^6,11,12,46,47^. The identification of LC2 as a cleavage-derived adaptor that links cargo to motors raises the possibility that similar mechanisms operate more widely to regulate protein localisation in neurons.

In summary, we propose that Calpain-10–mediated cleavage of MAP1A generates LC2, which links TRIM46 to kinesin-2 motors for delivery to the proximal axon. This mechanism couples proteolytic processing to intracellular transport and cytoskeletal organisation, thereby maintaining axonal structure in mature neurons and providing a new model for understanding how neuronal polarity and function is preserved.

## Acknowledgements

Microscopy was performed in the Otago Micro and Nanoscale Imaging Confocal Microscopy Unit, Research Infrastructure Centre at the University of Otago, Dunedin, New Zealand. We thank Graceallah Suhono for initial contributions to this project. We are grateful to Peter Fineran, Anastasia Labudina and Amy Jones for valuable feedback on this manuscript.

## Author contributions

MP: methodology, validation, formal analysis, investigation, writing – original draft, visualisation. EKG: methodology, validation, formal analysis, investigation, writing – original draft, visualisation. LFG: conceptualisation, resources, writing - original draft, supervision, project administration, funding acquisition.

## Funding statement

The authors disclose receipt of the following financial support for the research, authorship, and/or publication of this article: This work was supported by the Royal Society of New Zealand, Te Apārangi Marsden Fund [grant numbers 18-UOO-174 and 22-UOO-70] and Health Research Council of New Zealand [grant number 21-080].

---

**Supplementary Figure 1.**
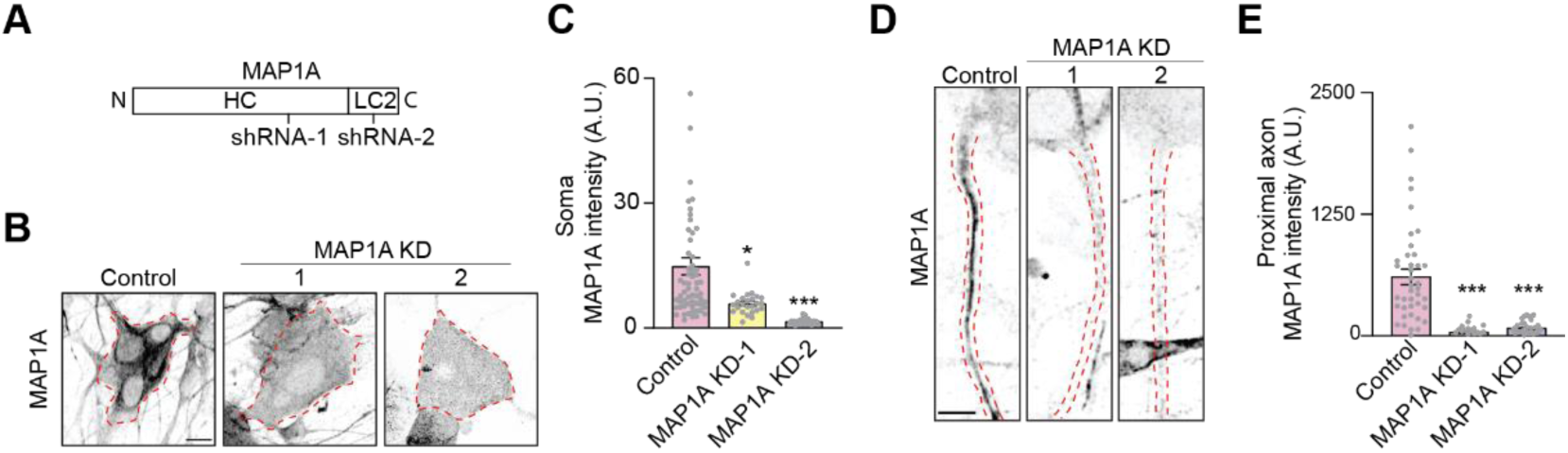
related to Figure 2. MAP1A-targeting shRNAs result in efficient knockdown of MAP1A expression. (A) Schematic showing the locations of shRNAs targeting MAP1A along the MAP1A mRNA sequence. (B) Representative images of the soma of adult sensory neurons (5 DIV) immunostained for endogenous MAP1A under the indicated conditions (KD=knockdown). Dashed red line outlines soma. (C) Average fluorescent intensity of MAP1A in the soma under the indicated conditions (control n=61, MAP1A KD-1 n=25, MAP1A KD-2 n=32). (D) Representative images of the proximal of adult sensory neurons (5 DIV) immunostained for endogenous MAP1A under the indicated conditions. Dashed red line highlights soma. (E) Average fluorescent intensity of MAP1A in the proximal axon under the indicated conditions (control n=40, MAP1A KD-1 n=16, MAP1A KD-2 n=28). n=neurons, A.U., arbitrary units. Mean ± SEM. Scale bars indicate 10 µm in (B) and (D). ***p <0.001; **p < 0.01; *p <0.05 Kruskal-Wallis test, with Dunn’s multiple comparisons post-hoc analysis.

**Supplementary Figure 2.**
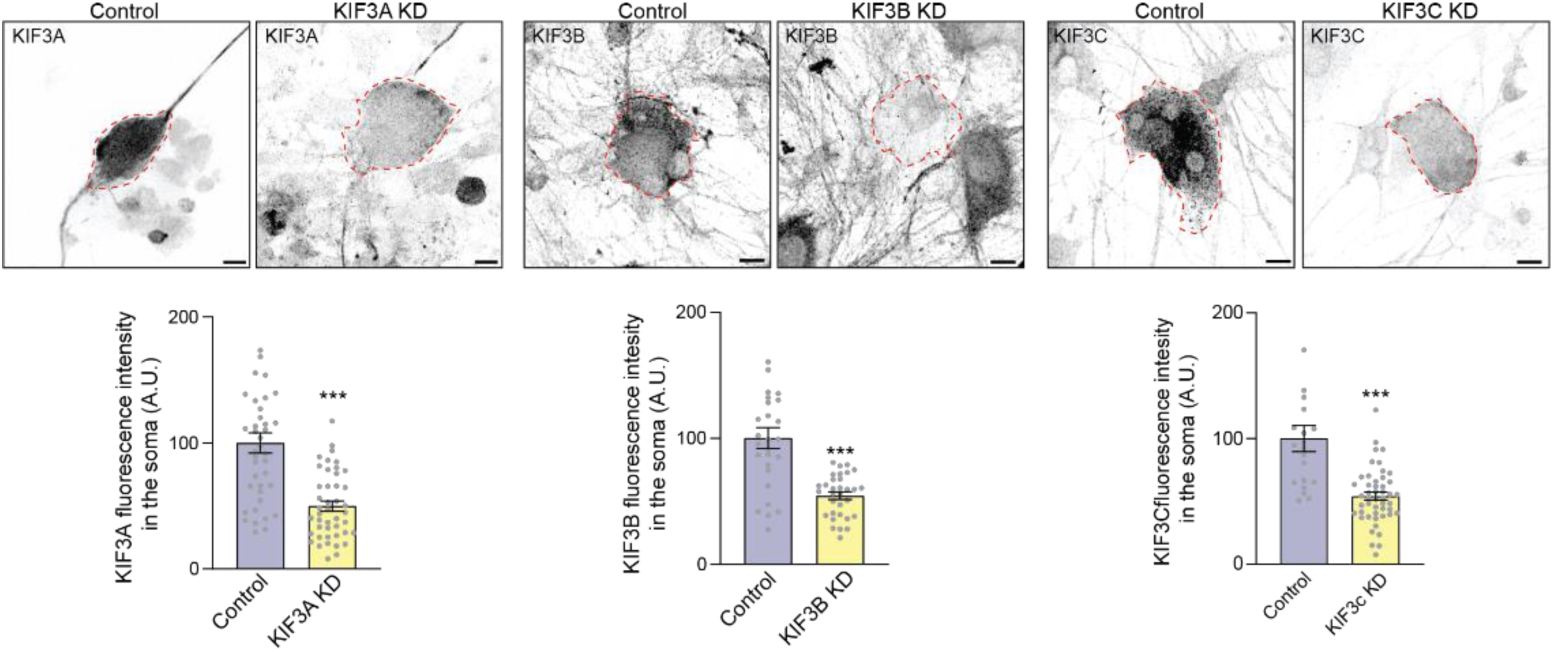
related to Figure 5. Validation of KIF3-targeting shRNAs. (A) Representative images of the soma of adult sensory neurons (5 DIV) immunostained for endogenous MAP1A under the indicated conditions (KD=knockdown). Dashed red line outlines soma of a transfected neuron. Graph shows average fluorescent intensity of KIF3A in the soma under the indicated conditions (control n=38, KIF3A KD=44). (B) Representative images of the soma of adult sensory neurons (5 DIV) immunostained for endogenous KIF3B under the indicated conditions (KD=knockdown). Dashed red line outlines soma of a transfected neuron. Graph shows average fluorescent intensity of KIF3B in the soma under the indicated conditions (control n=26, KIF3B KD=32). (C) Representative images of the soma of adult sensory neurons (5 DIV) immunostained for endogenous MAP1A under the indicated conditions (KD=knockdown). Dashed red line outlines soma of a transfected neuron. Graph shows average fluorescent intensity of KIF3C in the soma under the indicated conditions (control n=19, KIF3C KD=45). n=neurons, A.U., arbitrary units. Mean ± SEM. Scale bars indicate 10 µm. ***p <0.001; Kruskal-Wallis test, with Dunn’s multiple comparisons post-hoc analysis.

## Notes

### Competing Interest Statement

The authors have declared no competing interest.

